# Experimental heatwaves and warming cause distinctive community responses through their interactions with a novel species

**DOI:** 10.1101/2023.03.24.534073

**Authors:** Jinlin Chen, Owen T. Lewis

## Abstract

As mean temperatures increase and heatwaves become more frequent, species are expanding their distributions to colonise new habitats. The resulting novel species interactions will simultaneously shape the temperature-driven reorganization of resident communities. The interactive effects of climate change and climate change-facilitated invasion have rarely been studied in multi-trophic communities, and are likely to differ depending on the nature of the climatic driver (i.e. climate extremes or constant warming). We recreated under laboratory conditions a host-parasitoid community typical of high-elevation rainforest sites in Queensland, Australia, comprising four *Drosophila* species and two associated parasitoid species. We subjected these communities to climate change in the form of either heatwaves or constant warming, in combination with an invasion treatment involving a novel host species from lower-elevation habitats. The two parasitoid species were sensitive to both warming and heatwaves, while the demographic responses of *Drosophila* species were highly idiosyncratic, reflecting the combined effects of thermal tolerance, parasitism, competition, and facilitation. After multiple generations, heatwaves (but not constant warming) promoted the establishment of low-elevation species in upland communities. The introduction of this invading species correlated negatively with the abundance of one of the parasitoid species, leading to cascading effects on its hosts and their competitors. Our study, therefore, reveals differing, sometimes contrasting, impacts of extreme temperatures and constant warming on community composition. It also highlights how the scale and direction of climate impacts could be further modified by range-expanding species within a bi-trophic community network.

## Introduction

Climate change is leading to large-scale shifts in species’ distributions (Chen et al., 2011; Parmesan et al., 1999; Parmesan & Yohe, 2003). Differences in the speed (Mamantov et al., 2021), magnitude (Chen et al., 2011) and even direction (Lenoir et al., 2010) of range shifts are causing existing interactions among species to unravel, and new interactions to form.

Despite the widespread occurrence of such novel interactions, and evidence that they can influence community responses to climate change (Alexander, Diez, & Levine, 2015; Gilman, et al., 2010; Nomoto & Alexander, 2021), the ecological ramifications of climate-driven novel species interactions in multi-trophic communities are poorly explored. The impacts of climate change and novel species interactions on a biological community are mutually dependent (Figure 1). Climate change not only affects organisms’ performance but also alters the biotic environment (Blois, et al., 2013; Davis, et al., 2022; Descombes et al., 2020; Nowicki, et al., 2021; Romero et al., 2018). Climate-driven changes in the demography or behaviour of competitors, predators, and mutualists could modify the direct effects of climate on individual species. Such influential species can either be pre-existing components of the community (Barton & Ives 2014; Harmon, Moran, & Ives 2009) or novel species whose arrival is facilitated by climate change (Richman et al. 2020; Schweiger et al. 2010). Furthermore, species interactions, including those between novel species and residents, are themselves climate-dependent (Alexander et al., 2015; LaForgia, Harrison, & Latimer, 2020; Olsen, et al., 2016), as two of the dashed lines in Figure 1 represent. As a result, predicting biodiversity response to climate change is challenging because climate change leads to new species interactions as a result of range shifts and new climatic conditions to which species interactions are sensitive (Gilman et al., 2010; HilleRisLambers, et al., 2013; Tylianakis, et al., 2008; Wallingford et al., 2020).

**Figure 1.**
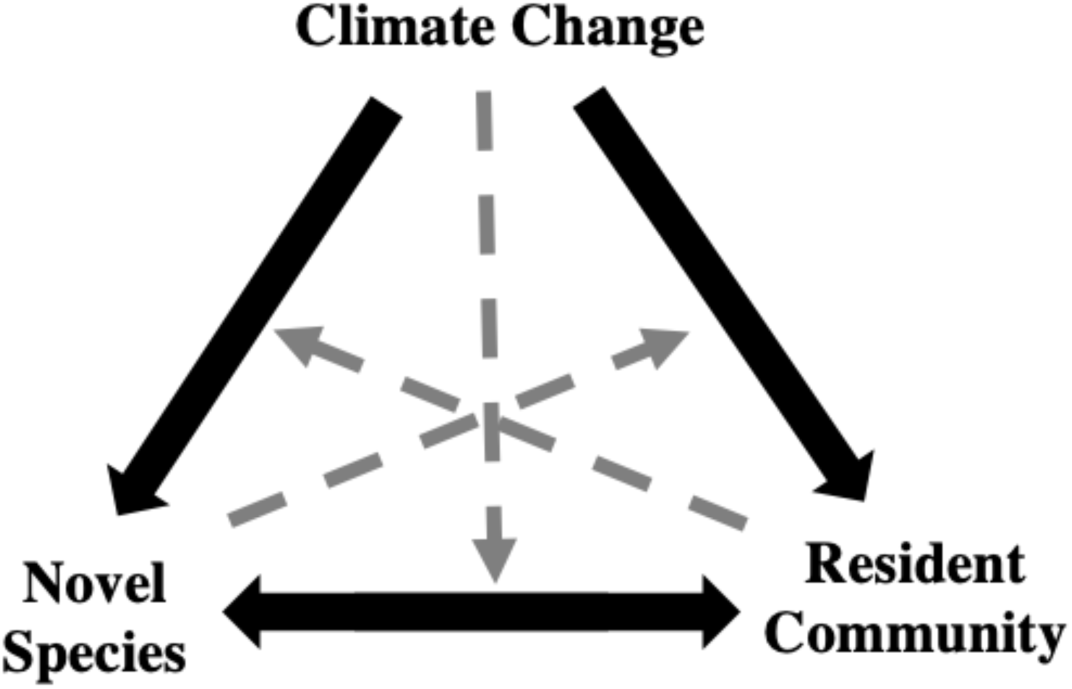
The relationships among climate change, novel species and the resident community. The solid arrows represent direct effects. Dashed arrows indicate that a third factor influences the direct effect between the two factors.

Experimental approaches have been useful for quantifying the impact of potential novel species interactions outside the current climatic conditions experienced by communities (Alexander, et al., 2016; Descombes et al., 2020; Ravi et al., 2022; Richman et al., 2020). Nevertheless, previous studies have usually focused on a single type of species interaction (e.g., competition, herbivory). Most biological communities, on the other hand, comprise at least a few trophic levels and several species within each level, resulting in motifs with a variety of trophic and non-trophic interaction pathways, both antagonistic and facilitative (Barton & Ives, 2014; Losapio et al., 2021; Thierry, Hrček, & Lewis, 2019). Understanding how the impacts of external drivers such as climate and invading species propagate through species interactions and emerge in multi-species, multi-trophic communities remains a key challenge.

While most empirical research on range shifts has focused on the impact of a constant increase of mean temperature (referred to here as “warming”), climate change also manifests as extreme high-temperature events and higher temperature variability, especially in tropical regions (IPCC, 2022; Perkins-Kirkpatrick & Lewis, 2020). Extreme events such as heatwaves also contribute to range dynamics (Diez et al., 2012; Terry, O’Sullivan, & Rossberg, 2022; Vasseur et al., 2014), prompting interest in the relative importance of warming versus extreme temperature events (Jentsch, Kreyling, & Beierkuhnlein, 2007; Lynch et al., 2014; G. Ma, Hoffmann, & Ma, 2015). A moderate level of warming generally increases interaction intensity (Klanderud, Vandvik, & Goldberg, 2015; Roslin et al., 2017) or releases physiological constraints (Crozier, 2004). In contrast, heatwaves often induce mortality or infertility (van Heerwaarden & Sgrò, 2021), temporarily releasing the surviving species from antagonistic biotic interactions (Diez et al., 2012; Wallingford et al., 2020). Different mitigation strategies are required for warming and heatwaves as they operate at different timescales. However, these strategies have seldom been put into practice because the ecological impacts of heatwaves versus warming have rarely been compared (C. Sen Ma, Ma, & Pincebourde, 2021).

Here, we investigate the interacting effects of novel species and different modes of climate change (i.e., warming versus heatwaves) on an experimental host-parasitoid system. Parasitoids are insects whose larvae develop on or in the bodies of other arthropods and eventually kill them (Hardy, Alphen, & Godfray, 1994). Host-parasitoid networks represent relatively tractable, strongly-interacting units containing a range of specialist and generalist interactions (Hrček & Godfray, 2015). Parasitoids play a major role in regulating the abundance and coexistence of arthropods in natural ecosystems and agriculture (Bonsall & Hassell, 1997; Hardy et al., 1994) and as such are often selected as bio-control agents. However, these ecological functions are especially vulnerable to climate change (Furlong & Zalucki, 2017; Hance, et al., 2007).

Our focal community comprises *Drosophila* species and their associated parasitoids from Australian rainforests (Jeffs et al., 2021). We focus on high-elevation communities as they are bounded geographically: as the climate warms, these communities face combined threats from changing abiotic and biotic conditions (Shah et al., 2020) without the presence of spatial refuges to buffer these threats. A previous study has shown that the upland community contains heat-sensitive species, and that interspecific competition prevents lowland species from occupying cooler, higher-elevation locations (J. Chen & Lewis, 2022). Based on this, we predict that heatwaves will have a larger effect than warming on community composition through their detrimental impact on heat-sensitive species. Second, we predict that heatwaves, but not warming, will facilitate the range expansion of a lowland species by releasing competitive pressure. Third, we predict that the impact of invasion will depend on climate scenarios as the low-elevation invader is only expected to successfully establish in the heatwaves treatment. Finally, we expect parasitoids to play a significant role in mediating the impact of temperature and/or the introduction of the novel species.

## Materials and Methods

### Study system

We conducted an experiment using tropical forest *Drosophila* flies and parasitoids in a controlled laboratory environment. Cultures of all species were collected from the Wet Tropics bioregion in Queensland, Australia, which has a high level of species endemism associated with cool, moist upland refugia. Information on the natural distribution patterns of *Drosophila* and their natural enemies, parasitoids, is based on previous trapping using banana baits (Jeffs et al., 2021). Most *Drosophila* species and some of the parasitoids from these communities are now in laboratory culture, allowing multi-species manipulative experiments in the laboratory.

We assembled an experimental community to reflect species compositions at contemporary high-elevation sites (elevation 730m – 880m), comprising *D. birchii*, *D. rubida*, *D. pseudotakahashii*, *D. pallidifrons*, a generalist pupal parasitoid from the genus *Trichopria* (lab strain 66LD, species identifier drop-Dia127-sp9), and a more specialized larval parasitoid from the genus *Asobara* (lab strain KHB, species identifier drop-Aso-sp8) (Lue et al., 2021). The species used and the host specificity of the parasitoids (Chia-Hua Lue, personal communication) are shown in Figure 2a. The four *Drosophila* species account for 77% – 95% of *Drosophila* abundance sampled at high-elevation sites. The distribution of the two parasitoid species is unclear because of their small sample sizes, but both occur at high-elevation sites (Jeffs et al., 2021). To represent a lowland species which is a candidate to invade higher-elevation communities under climate change, we used *D. pandora*. It occurs predominantly at low-elevation (70m) sites (Jeffs et al. 2021) and is known from thermal performance trials to be adapted to higher temperatures (J. Chen and Lewis, 2022).

**Figure 2.**
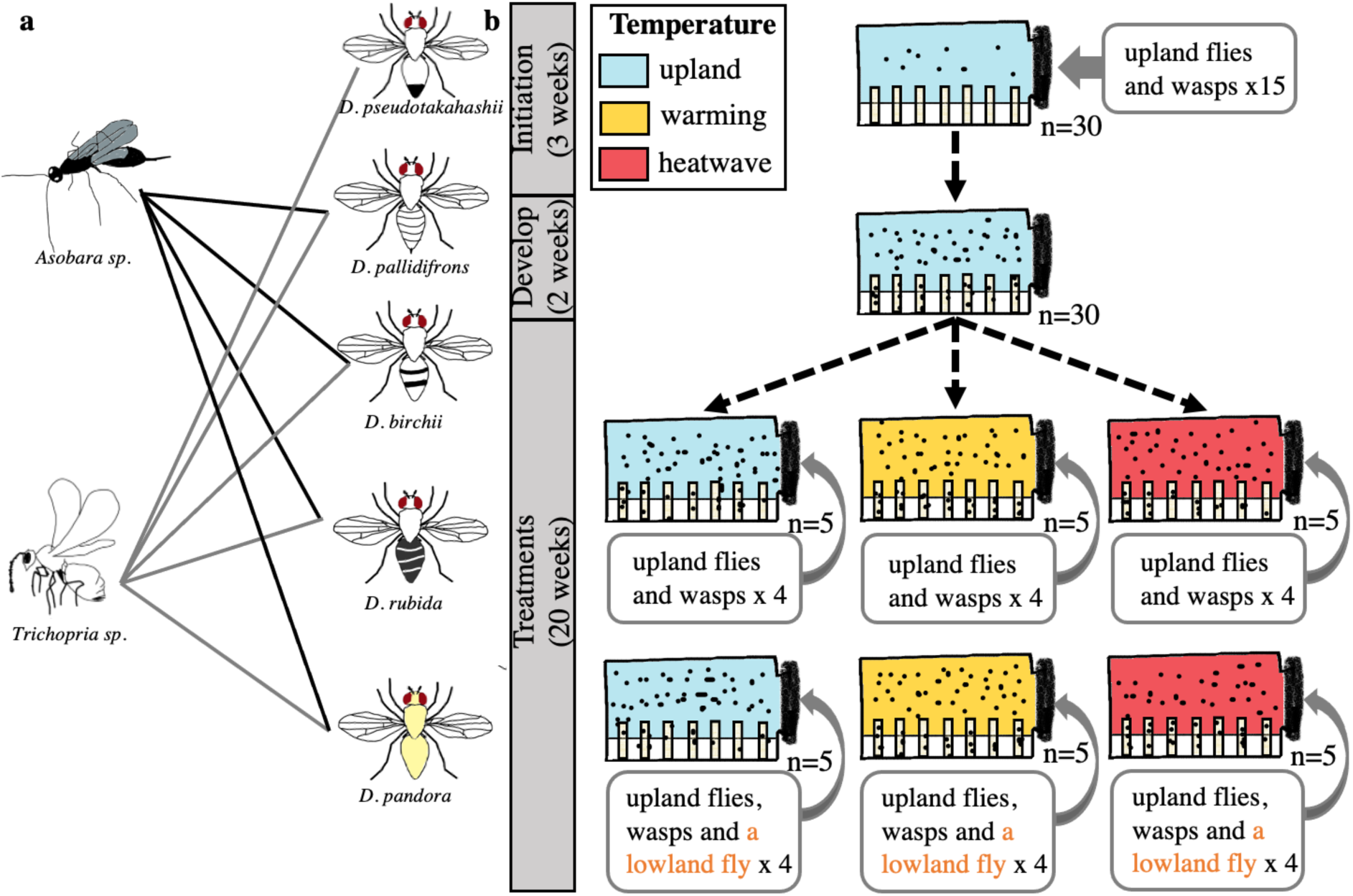
The study system and experimental procedures. (a) The known trophic interactions between parasitoids and *Drosophila* flies, based on observation of the successful development of the parasitoids. *Drosophila pandora* is a low-elevation specialist, used as a species that is novel to the upland community. Different *Drosophila* species are illustrated by differing abdomen patterns, and these drawings are used in subsequent figures to represent the corresponding species. (b) Experimental communities were maintained in sealed cages with regular food provisions. Fifteen pairs of each upland species were added to every cage for the first three weeks. Communities then developed for two additional weeks. Following this, a total of 30 cages (five replicates per treatment combination) underwent combinations of novel species and temperature treatments (blue: upland temperature; yellow: warming; red: heatwave). The boxed text under each cage indicates the species, and the number of pairs of each species, added to the cage each week.

### Community initiation

All experimental communities were initiated with the same species on a day/night cycle (24°C, light for 12h; 18°C, dark for 12h), mimicking high-elevation conditions (also referred to as upland or control conditions). Each community was kept in a sealed 20cm x 5cm x 10cm plastic cage. Card dividers on the floor of the cage could hold up to 24 standard *Drosophila* vials. In each of the first three weeks, 15 females and 15 males of *D. pseudotakahashii*, *D. palidifrons*, *D. birchii*, *D. rubida,* and 10 females and 10 males of both parasitoid species were added to each cage. The adult flies were of mixed ages, randomly sourced from mass-bred population cages. The parasitoids were one-week-old adults, reared on *D. melanogaster*, a species that does not exist in our study site. After these first three weeks, we maintained the community cages without adding insects for two more weeks before starting experimental treatments. The schedule is shown in Figure 2b.

### Maintenance and treatments

After the fifth week of community initiation, a factorial design of three temperature treatments and two introduction treatments was applied to 30 cages (18 cages in block A and 12 cages in block B, initiated four weeks apart). Five cages were randomly assigned to each treatment and maintained for another 20 weeks. Three vials of 6ml fresh *Drosophila* food (weight/volume concentration: 8% corn flour, 4% yeast, 5% sugar, 1% agar, and 1.67% methyl-4-hydroxybenzoate) were added to the cage twice a week (three to four days apart). Vials added in the middle of each week were “maintenance vials”, which were removed after 40 days (most flies and parasitoids had emerged by then). Vials added at the start of the week were “sampling vials”, which were removed after ten days, before the emergence of any insects from the vials.

#### Temperature regimes

As described above, the upland temperature regime alternated between 24°C, light (12h) and 18°C, dark (12h). For warming and heatwave treatments, we allocated the same degree-day increase in cumulative temperature (+1°C per day), either in the form of an increase in mean temperature or in the form of heatwave events. The warming regime increased both the day-time and night-time temperatures by 1°C (25°C, light for 12h; 19°C, dark for 12h), which is anticipated by the end of this century under the lowest emission scheme (IPCC, 2022). The heatwave regime had the same baseline temperature as the upland regime, but with periodic heatwaves when daytime and night-time temperatures rose by 6°C for five days. For comparison, in February 2017, a heatwave raised the maximum air temperature by 6°C and the average air temperature by 4°C for two consecutive days at our study site (Supplementary Figure 1a). Heatwaves were applied to all corresponding communities at the same time following a predetermined timing. Heatwaves occurred once every four weeks on average, with a two-week minimum interval between consecutive heatwaves (sequences of heatwave events are shown in Supplementary Figure 1b). We used three incubators of two similar models (Sanyo MIR-154 & Sanyo MIR-153). To avoid any confounding incubator effects, temperature regimes were rotated among incubators weekly.

#### Introductio

Every week, four pairs of the low-elevation specialist, *D. pandora*, were added to cages assigned for the introduction treatment. To avoid the stochastic extinction of resident species, and to mimic natural immigration, four pairs of each *Drosophila* and parasitoid species from the starting community were added to all community cages at the same frequency as *D. pandora*. In the final two weeks of the experiment, no animals were added.

### Data collection

#### Regular sampling

The reproductive success of every species was monitored by examining the adults emerging from the regularly sampled vials. After allowing ten days of colonization, the sampling vials were removed from cages and kept for 24 days in the same environmental regime as the communities from which they originated. Thus, the number of emerging adults from the three “sampling vials” reflects the combined impact of adult population sizes at sampling, adult fecundity, intra- and inter-specific competition, and parasitism. Emerging fly and parasitoid adults were collected every three or four days and stored by freezing at −18°C. Adults were identified to species, sexed and counted. Weekly counts of the three sampling vials from the same community were summed.

#### Heatwave-specific sampling

To measure how a single heatwave event altered the reproductive success of flies, fresh food vials were placed into each cage at three time points relative to the final heatwave event: one day before the final heatwave, on the fifth day of the heatwave, and five days after the heatwave ended. At each time point, three vials (each containing 6ml of food) were added to each relevant cage for 24 hours, a sufficiently short exposure to avoid larval crowding within vials. These vials were then kept under the upland temperature regime and emerging adult flies were identified and counted. There were no parasitoid offspring as these parasitoids cannot parasitise eggs or first-instar larvae.

#### Termination and census

After 25 weeks, all living adult insects in community cages were frozen, identified and counted. Throughout the maintenance of the community cages, the numbers of *Drosophila* or parasitoids increased but did not appear to fluctuate greatly. Therefore, although the final census is only a snapshot we regard it as a useful representation of community composition. Females of *D. pandora*, *D. pseudotakahashii*, and *D. birchii* are marginally different in pigmentation and body size, but become difficult to separate in crowded environments which affect their body sizes. Analyses of these three species were therefore based on counts of males if not otherwise specified.

### Data analysis

All statistical analyses were performed with R version 4.0.3 (R Core Team, 2020). Following the fitting of (generalised) linear models, diagnostic checks of residuals highlighted no issues.

#### Community composition

Community compositions of the living adults at the end of the maintenance period were analysed separately for hosts and parasitoids. Compositions of four resident *Drosophila* were visualized using non-metric multidimensional scaling (NMDs) with a dimension of two. Abundances of parasitoids were standardized as z-scores (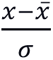). Community distance matrices were calculated using the “bray” method (9999 permutations) and assessed through non-parametric MANOVA (Anderson, 2001). Upon detecting overall significant differences among all treatments, post-hoc analyses were performed to test the differences between pairs of treatments. NMDs and non-parametric MANOVA analyses were conducted using the *vegan* package (Oksanen, et al., 2020). The *multcompView* package was used to summarise significant pairwise differences based on post-hoc comparisons (Graves, Piepho & Selzer, 2019).

Species responses to treatments were then analysed individually using generalized linear models. The census abundance of each resident species was modelled as a function of *temperature treatment*, *the introduction of the novel species*, their interaction term, and *block,* assuming Poisson error distributions. P values of these predictors of all upland species were adjusted by Bonferroni correction. Post-hoc pairwise comparisons were conducted using the *emmeans* package (Lenth, 2022).

#### Effect of temperature treatment on the introduced species

The final adult abundance of *D. pandora* was modelled as a function of *temperature treatment*, *block* and their interaction term in a generalized linear model with Poisson error distribution. For tri-weekly sampling data, *D. pandora* numbers were modelled as a function of *temperature treatment* and *block* as fixed effects, *week of sampling,* and *cage ID* as random effects in a generalized linear mixed-effect model with Poisson errors.

#### Effect of heatwave treatment on reproduction of Drosophila species

Counts were log-transformed (log(y+1)) and then modelled as a function of *sampling time* (relative to the heatwave event), *species*, *the introduction of the novel species*, *block*, and the interaction between *sampling time* and *species* in a linear model. This strategy was used because residuals were not homogeneous when counts were modelled directly, assuming Poisson errors, in a generalized linear mixed-effect model. The Linear mixed-effect model used the *lme4* package (Bates, et al., 2015).

#### Inferring indirect effects

To examine whether the impact of a treatment on a species is direct or mediated via other species, we tested whether the effect coefficient of a treatment was reduced by the inclusion in the regression model of the putative mediating species. Directed Acyclic Graphs (DAGs) were built to describe the direct and potential indirect pathways by which temperature and introduction treatments affect the reproductive outputs of three focal *Drosophila*: *D. pandora*, *D. pallidifrons* and *D. pseudotakahashii*. In the DAGs, the two parasitoids were the potential mediators between all treatments and all focal species. Additionally, the known dominant competitor (J. Chen & Lewis, 2022), *D. pallidifrons*, was hypothesized to affect adult offspring numbers of *D. pandora* and *D. pseudotakahashii* directly by competition.

We first built regression models to identify the effects of treatments on the reproductive success of parasitoids. In these generalized linear mixed-effect models (with Poisson error distributions), we used *temperature treatment*, *the introduction of the novel species*, their interaction, and *block* as fixed effects, and *sampling week* and *cage ID* as random effects. P values of all tests on parasitoids were then adjusted by Bonferroni correction. For each focal *Drosophila* species, we first build a regression model following the same modelling strategy as for parasitoids, to test for the overall effects of the experimental treatments. If treatments had significant effects, a further regression model was constructed based on the DAG (Figure 6, left panel), adding standardized numbers (z-scores) of parasitoids and/or other relevant *Drosophila* as additional predictors. P values from the above two regressions of each *Drosophila* species were grouped and adjusted for multiple comparisons. Model outputs were then compared. If the fitted coefficient of the treatment was reduced after a significant controlling factor was added, we inferred that the overall effect of that treatment was (partially) mediated by the mediating species. The coefficients of treatments in the second model were direct effects which were not mediated by other species.

## Results

### Interacting effects of temperature and invaders on the resident community

The temperature treatments and the introduction of the novel species jointly caused compositional changes in upland communities. Non-parametric MANOVA identified significant differences among treatments in the community composition of upland *Drosophila* and parasitoid species (*Drosophila*: F_5,24_ = 7.8, p = 0.0001, r^2^ = 0.62; parasitoids: F_5,24_ = 6.5, p = 0.0003, r^2^ = 0.58). Post-hoc pairwise analysis revealed significant differences (denoted by different letters on polygons in Figure 3) between groups of communities. Details on individual species are shown in Supplemental Figure 2. Treatment effects on individual species were analysed using regression models, shown in Supplementary Table 1.

**Figure 3.**
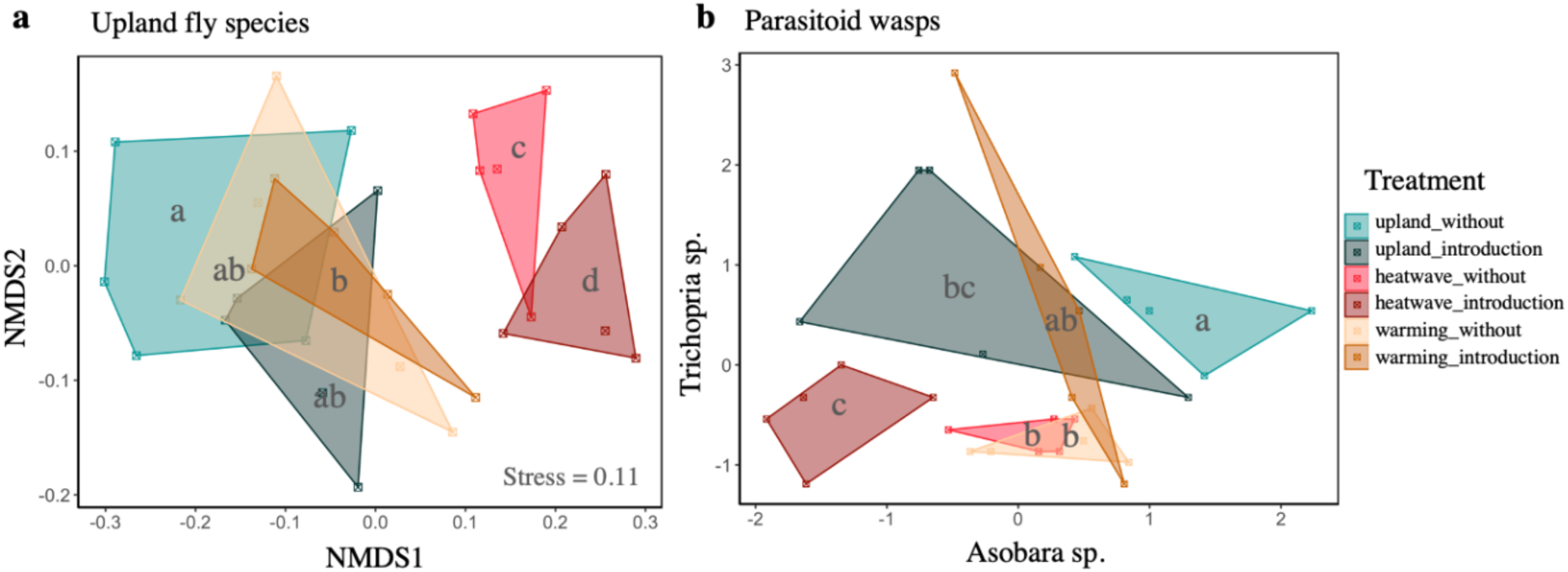
Compositional changes of hosts (a) and parasitoids (b) under different combinations of temperature and introduction treatments. Hosts composition (four *Drosophila* species) is visualized in two dimensions using NMDS. For parasitoids, their standardized abundances are shown on the axes. Each point represents a community. Communities of the same treatment are grouped by a minimal convex polygon (replicate=5). The colours of polygons correspond to treatments, as described in the panel. Letters inside the polygons indicate statistically different groups tested by non-parametric MANOVA.

Heatwaves, but not warming, significantly changed the community composition of resident *Drosophila* (Figure 3a). This change was driven by a large decrease in *D. pseudotakahashii* and a small increase in *pallidifrons* (Supplementary Table 1 and Supplementary Figure 2). Introducing the novel species only had a significant effect on the local *Drosophila* community when concurrent with the heatwave treatment. Heatwaves and invasion jointly resulted in communities dominated by *D. pallidifrons.* The abundances of *D. birchii* and *D. rubida* stay low across treatments (Supplementary Figures 2e and f).

Warming and heatwaves reduced the relative abundances of both parasitoid species by a similar degree in the absence of the novel species (Figure 3b). Introducing the novel species significantly changed the parasitoid composition by reducing the abundance of *Asobara* parasitoids under both the upland temperature and heatwave treatment, but had no detectable effect under warming conditions.

#### Performance of invading species in different temperature treatments

At the end of the experiment, the censused abundances of the invader *D. pandora* in the upland temperature (control) and warming treatments were both low (Figure 4a; p = 0.75; degree of freedom = 9). From the same model, the abundance of *D. pandora* in heatwave treatments was significantly higher (Figure 4a; coefficient = 2.02, p < 0.001). The heatwave treatment still had a positive effect in the second block, although smaller (coefficient of heatwave x block = −0.67, p = 0.008). Consistent with the census, *D. pandora* produced significantly higher numbers of adult offspring in the heatwave treatment (Figure 4b; coefficient of heatwave treatment = 2.12, p < 0.0001). The effect of warming was not significant compared with the upland control (p = 0.73). Very few offspring were produced in either upland temperature or warming treatment. The second block had fewer offspring than the first (coefficient of block = −0.44, p = 0.04).

**Figure 4.**
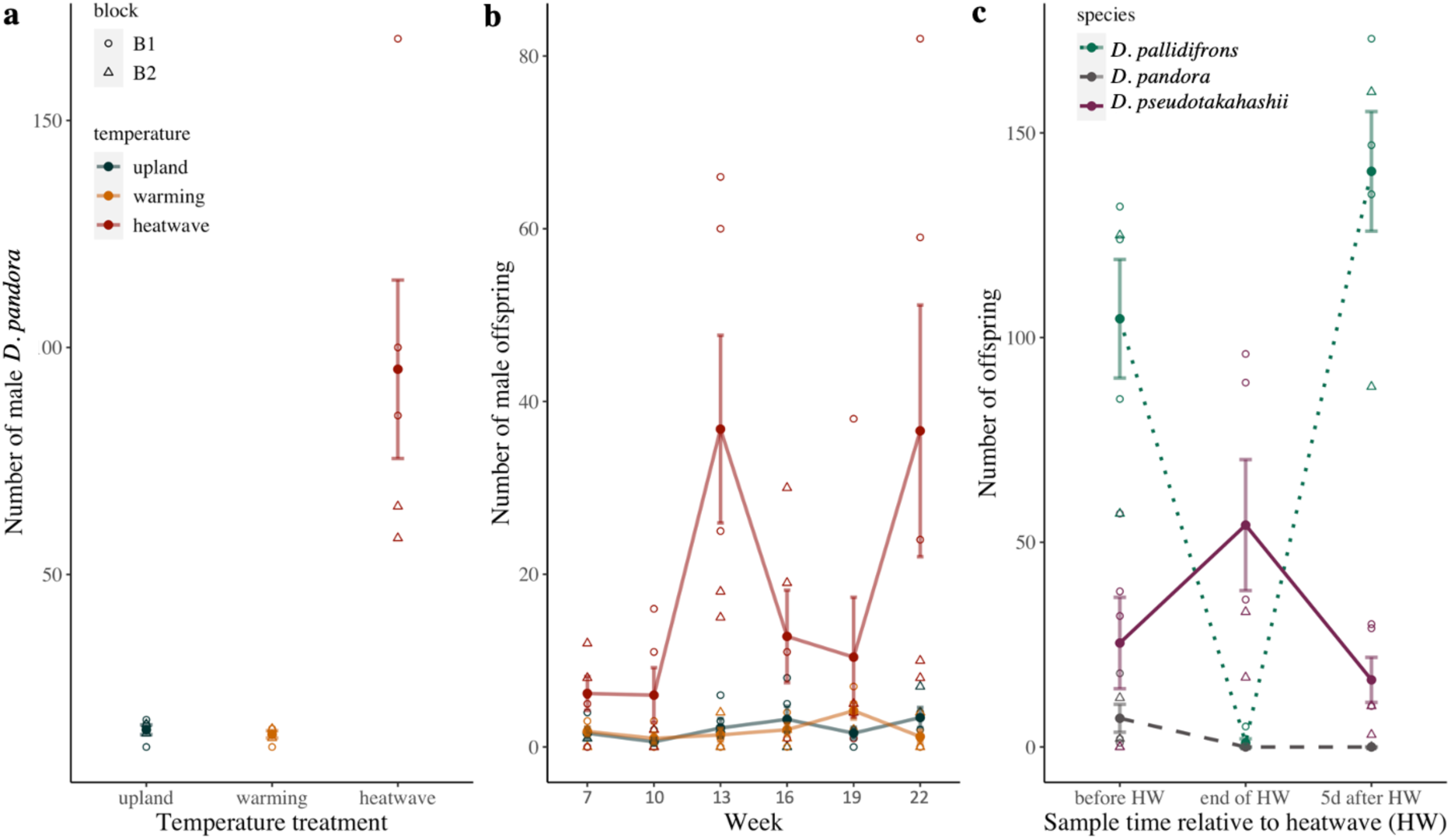
Heatwaves promote the establishment of the lowland specialist, *D. pandora*, in upland communities. (a) Numbers of male *D. pandora* adults in community cages upon termination under different temperature treatments. (b) Reproduction of *D. pandora* over time in the upland temperature regime (blue), warming treatment (yellow) and heatwave treatment (red). (c) Reproductive success of *D. pallidifrons* (green, dotted line), *D. pseudotakahashii* (grey, dashed line), and *D. pandora* (purple, solid line) on a single day at different timings relative to the last heatwave event. For all graphs, each open symbol represents the number of individuals censused or sampled from a community replicate (n = 5). The solid circles represent the means of the treatment groups. Shapes of symbols represent experimental block 1 (circle) or block 2 (triangle). Error bars show ±1 SD.

To investigate how heatwaves favoured *D. pandora*, we compared one-day snapshots of reproductive success at different timings relative to the final heatwave event (Figure 4c). The heatwave significantly decreased the reproductive success of *D. pallidifrons* (coefficient = −3.9, p < 0.001, degree of freedom = 64), but its reproduction recovered to a similar level as before the heatwave (p = 0.14). The heatwave also significantly decreased the reproductive success of *D. pseudotakahashii* to zero (p < 0.001), and it was unable to resume fertility within at least five days. In contrast, the heatwave temporarily increased the reproductive success of the invader *D. pandora* (coefficient = 1.57, p = 0.002).

### Parasitoid-mediated effects of treatments

We used reproductive success over time (raw data are shown in Supplementary Figure 3) to explore whether the effects of treatments on *D. pandora*, *D. pallidifrons* and *D. pseudotakahashii* were mediated by parasitoids’ top-down control. As shown in Table 1, the reproductive performance of *Asobara* was significantly reduced by invasion. Warming and heatwave treatments tend to have opposing effects on both *Asobara* and *Trichopria*. For the three focal *Drosophila* species, paths of treatment effects are visualized in Figure 5, and Table 2 shows the corresponding statistical analyses.

**Figure 5.**
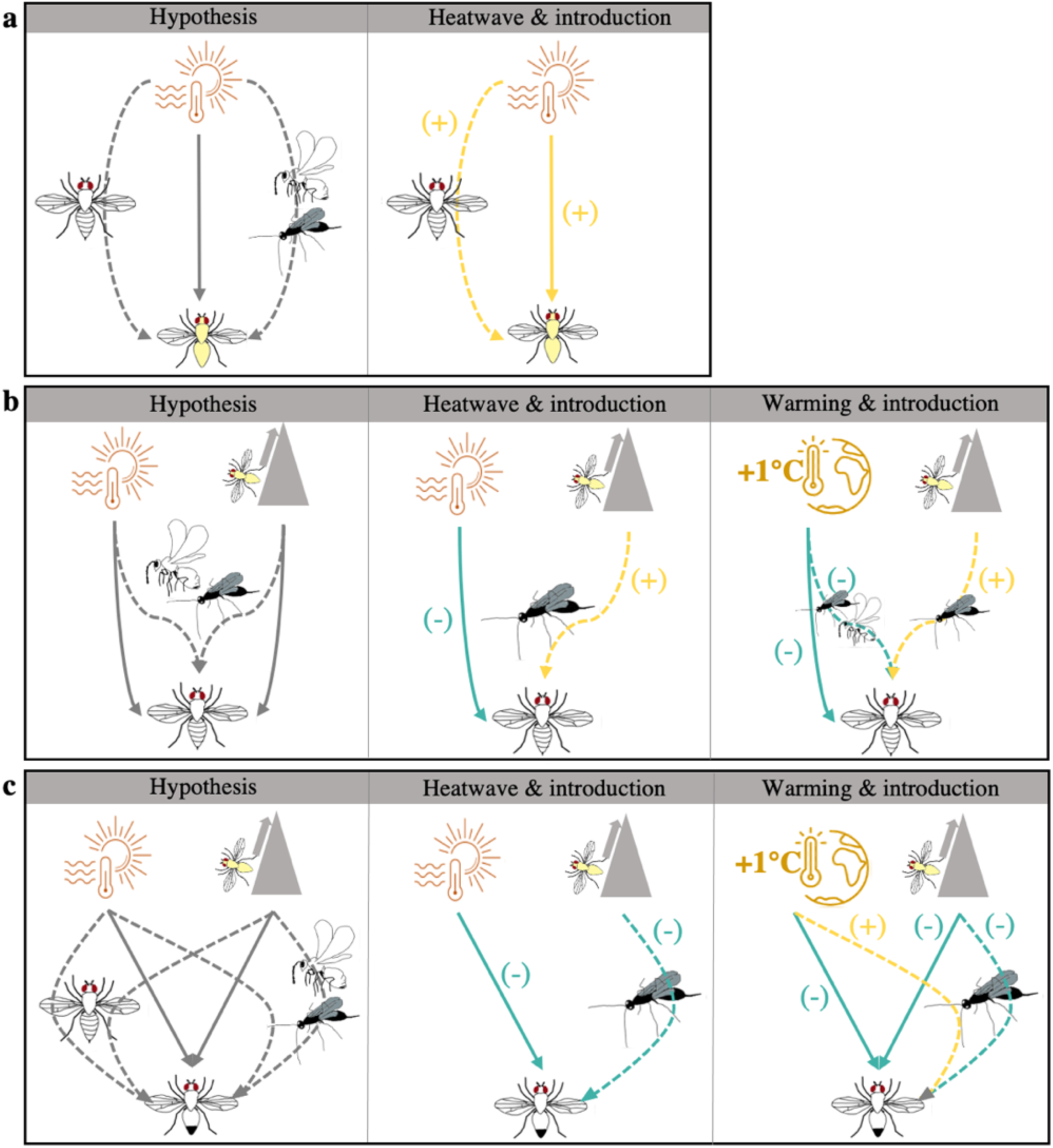
The direct and indirect effects of experimental treatments on the reproductive success of (a) *D. pandora*, (b) *D. pallidifrons* and (c) *D. pseudotakahashii*. The left panel shows DAGs which describe the hypothesized direct (solid line) and indirect (dashed lines) effects of experimental treatments on the focal species. The species symbols (defined in Figure 2) on top of the dashed lines represent the species mediating indirect effects. The middle column shows the effects of heatwave and introduction treatment for each *Drosophila* species. The right column shows the effects of warming and introduction treatment. Blue arrows indicate negative (-) effects and yellow arrows indicate positive (+) effects. Bonferroni correction removed statistical significance only for the effect of invasion on *D. pseudotakahashii* in the middle panel of (c).

**Table 1.**
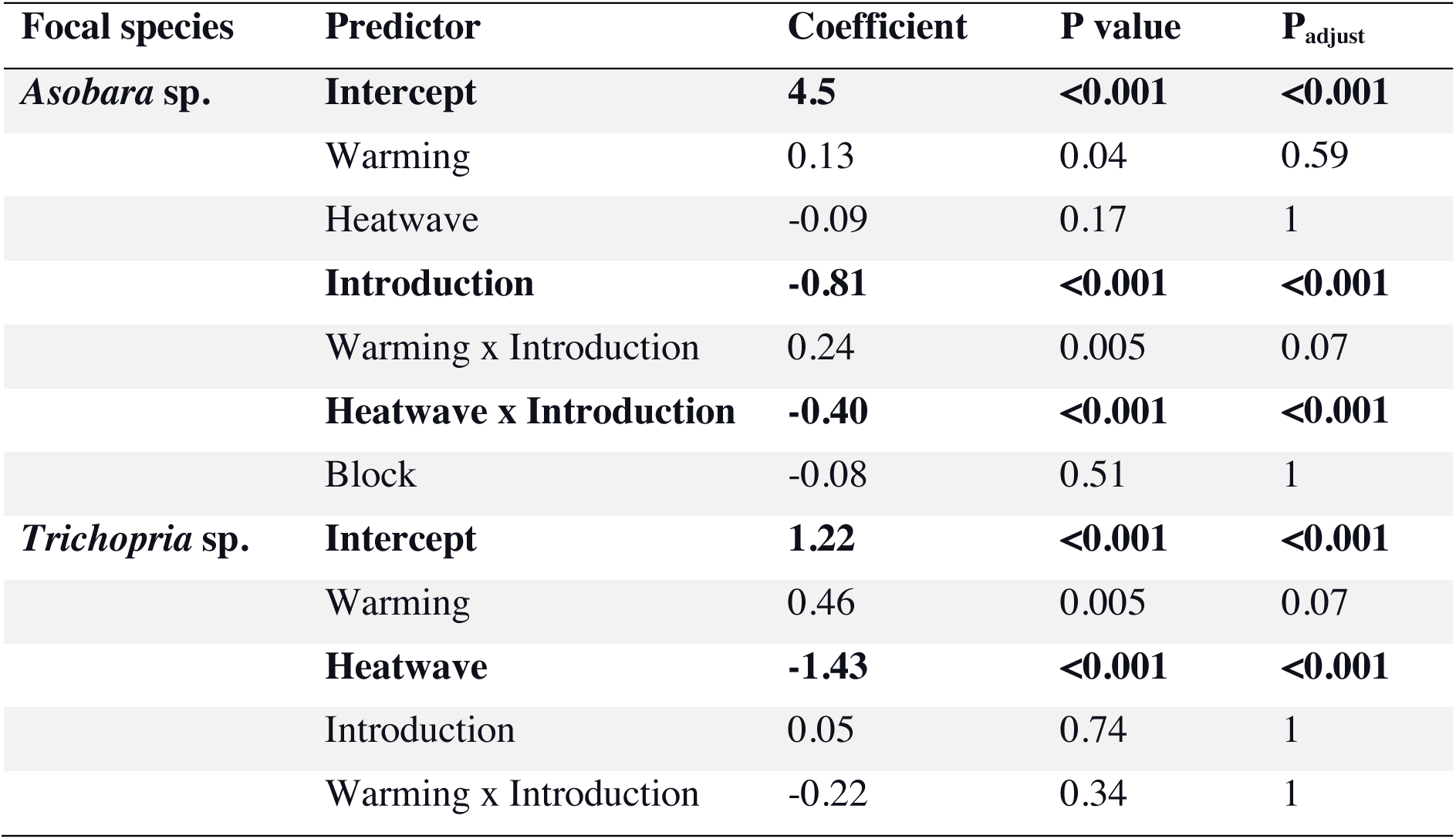
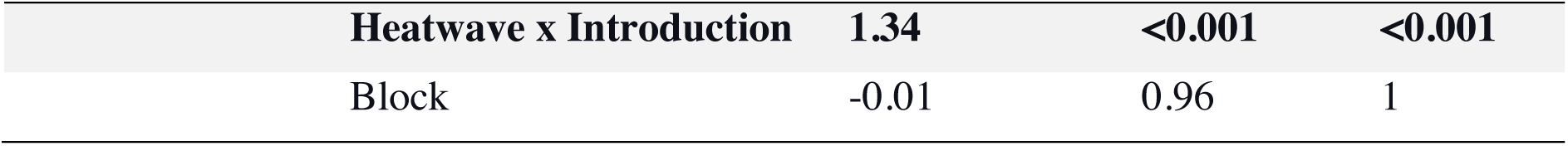
The effects of treatments and their interactions on the reproductive success of parasitoids. Numbers of individual species are analysed on a log scale. P_adjusted_ is the adjusted p value following Bonferroni correction. Significant effects after correction for multiple comparisons are in bold.

**Table 2.**
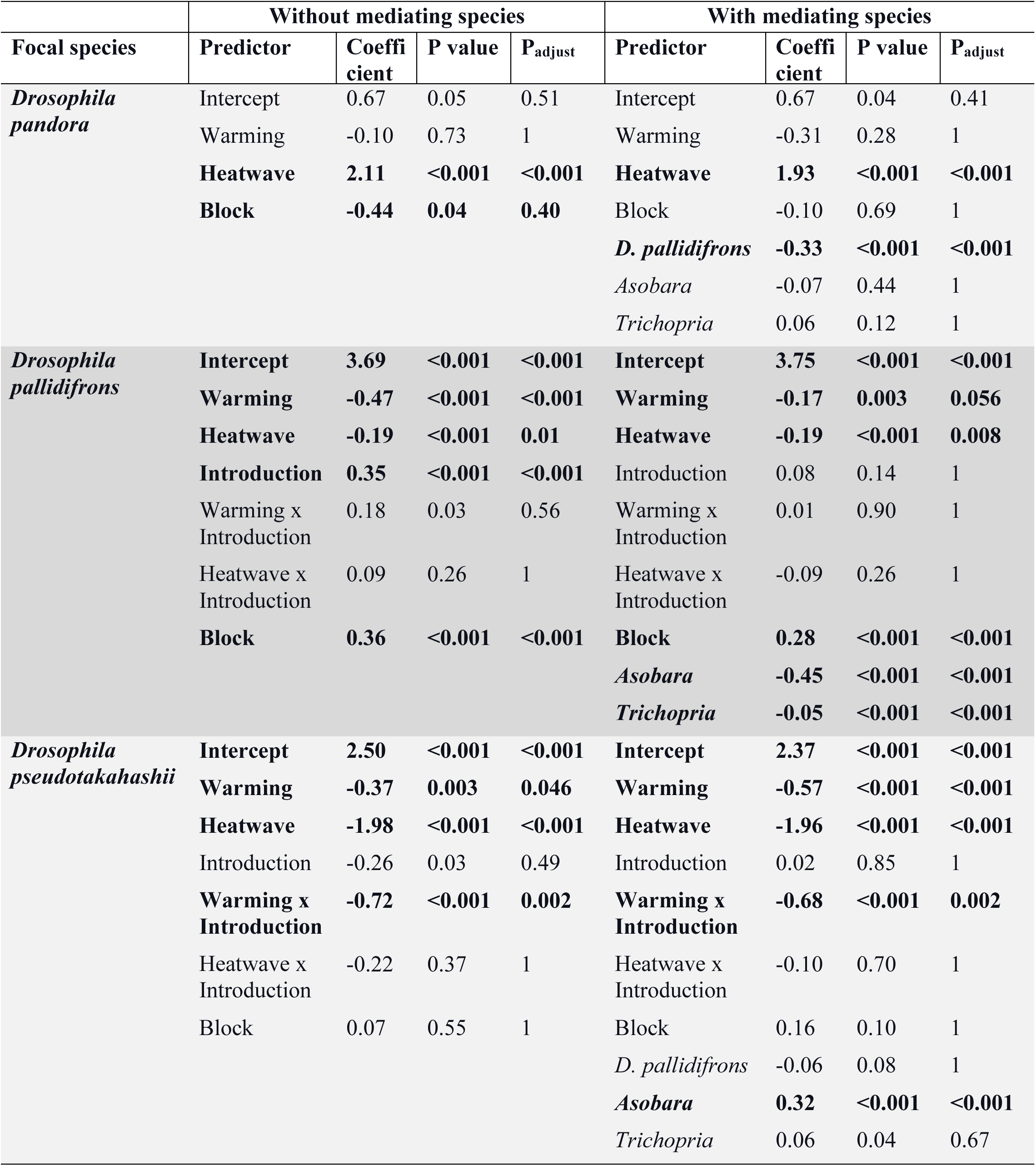
Regression testing the effects of experimental treatments on reproduction of the focal species, with and without controlling for mediating species. Numbers of individual species were log-transformed for analyses. P_adjusted_ is the adjusted p value following Bonferroni correction. Statistically significant effects after correction for multiple comparisons are printed in bold.

For *D. pandora*, controlling for the observed reproductive success of putative mediating species slightly decreased the coefficient of the heatwave treatment (coefficients before vs. after controlling for other species: 2.12 (p_adjust_ < 0.001) vs. 1.93 (p_adjust_ < 0.001)). The number of *D. pallidifrons* negatively correlated with the number of *D. pandora* (coefficient = −0.33, p_adjust_ < 0.0001), reflecting competition. The effects of both parasitoid species were non-significant.

Without controlling for other species, warming and heatwave both had significant negative effects on the reproduction of *D. pallidifrons*. Introducing the novel species significantly increased the reproduction of *D. pallidifrons*. When scaled numbers of parasitoids were added to the model, both parasitoid species had significant negative impacts. The effect of heatwave treatment was not changed (coefficients before vs. after: −0.19 (p_adjust_ = 0.01) vs. −0.19 (p_adjust_ = 0.008)). Warming still had a significant effect, but the effect was reduced (coefficients before vs. after: −0.47 (p_adjust_ < 0.001) vs. −0.17 (p_adjust_ = 0.056)), implying that the negative effect of warming is partly mediated by the positive effect of warming on parasitoids. The direct effect of introducing the invader was not significant after controlling for parasitoid number (p = 0.14), meaning that the positive effect of the invader on *D. pallidifrons* was completely mediated by parasitoids. The associations between both parasitoid species and *D. pallidifrons* were negative, reflecting parasitism.

For *D. pseudotakahashii*, warming and heatwave treatments had significant negative effects overall. Introducing the novel species significantly reduced *D. pseudotakahashii* reproduction under the warming treatment, but the effects of introduction were more marginal under other temperature treatments, becoming non-significant following Bonferroni correction. After controlling for the numbers of parasitoids and *D. pallidifrons*, the direct effect of warming was more negative than its overall effect (coefficients before vs. after: −0.37 (p_adjust_ = 0.046) vs. −0.57 (p_adjust_ < 0.001)). The negative effect of heatwaves remained the same after including mediating species in the models (coefficients before vs. after: −1.98 (p_adjust_ < 0.001) vs. −1.96 (p_adjust_ < 0.001)), implying that the effect of heatwaves on *D. pseudotakahashii* was mostly direct. Controlling for the numbers of mediating species reduced the coefficient of invasion on *D. pseudotakahashii*. The association between *Asobara* parasitoids and *D. pseudotakahashii* was significantly positive.

## Discussion

Our experimental community corresponds closely to the natural community of closely interacting *Drosophila* and parasitoids in the high-elevation rainforest of the Wet Tropics, Australia. We found that this community of up to seven interacting species in two trophic levels can respond in complex and unpredictable ways to different components of climate change and climate change-facilitated invasion, mediated by direct and indirect effects within the interaction network. Community structure changed more in response to periodic heatwaves than to the constant increase in mean temperature. Heatwaves, but not constant warming, also facilitated the establishment of a novel species associated with warmer, low-elevation forests. The invasion of this species additionally altered the composition of the wider community. Surprisingly, the invasion treatment significantly reduced the single-generation reproductive success and abundance of a larval parasitoid, *Asobara*, even when the novel species did not establish. Lastly, our data allow us to unravel potential mechanisms underlying these shifts in community composition and highlight the key role of cascading effects mediated by parasitoids within this relatively simple bi-trophic network.

### Effects of heatwaves and warming on the resident community

Threats to populations and communities from global warming depend on the mode of thermal stress (e.g., heatwaves or warming) and thermal sensitivities, which often vary among species or guilds (Furlong & Zalucki, 2017; G. Ma, Rudolf, & Ma, 2015). In our study, the heatwave treatment caused greater structural change than warming in the local *Drosophila* community. These results are consistent with previous studies which found that repeated, short, intense thermal stresses can cause more profound structural changes in communities than longer but less intense stresses (Polazzo, et al., 2023).

Analyses of individual species within these communities reveal that the impact of warming and heatwave treatments were different in magnitude and, sometimes, in direction. *D. pseudotakahashii* is the most heat-sensitive species in our study system (J. Chen and Lewis 2022). Its reproductive success and abundance were decreased by warming, and more so by heatwaves treatment. While heatwaves temporarily reduced the reproduction of *D. pallidifrons*, this species was able to resume reproduction quickly. Its high abundance in the heatwave treatment likely resulted from the long-term reduction of parasitoid numbers under this thermal regime. Warming unexpectedly decreased the single-generation reproduction and abundance of *D. pallidifrons*, perhaps because warming increased its fecundity, leading to larval and pupal crowding that greatly reduced their survival to adulthood (Ménsua and Moya 1983). The abundances of the two parasitoid species decreased to similar degrees under warming and heatwave treatments, although the direct effect of warming on their reproduction was positive, while heatwaves decreased their reproductive performance.

### Extreme heat events facilitate range expansion

Our results support the hypothesis that extreme heat events facilitate range expansion in our study system. Based on the asymmetrical shape of thermal performance curves, theories predict that the temporal distribution of rising temperatures is critical for organismal performance (Dell, Pawar, & Savage, 2014; Vasseur et al., 2014). We found that, with the same rise in mean temperature, periodic heatwaves had a stronger direct negative impact than constant warming on the single-generation reproductive success of the resident flies. This large decrease in the reproductive performance of the resident species reduced the biotic resistance of the dominant competitors, creating a temporal window when the novel species, *D. pandora*, was able to build up its numbers (Figure 6a).

**Figure 6.**
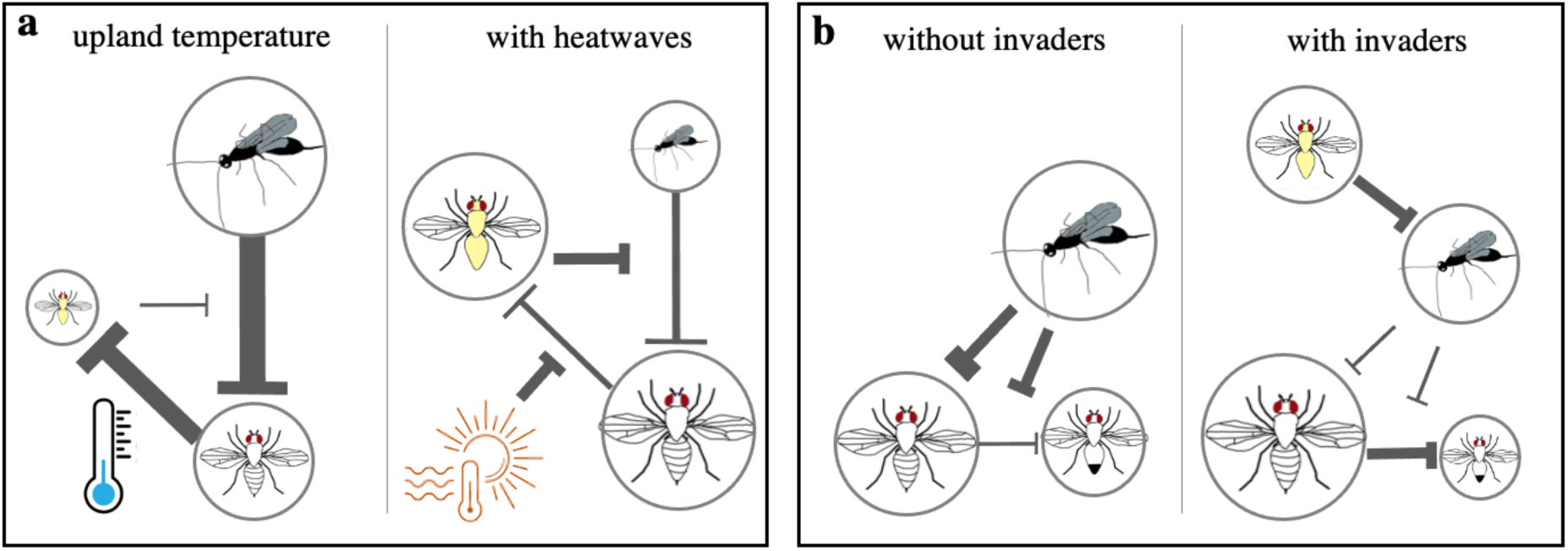
Key species interaction motifs that contribute to the community responses to treatments. (a) Summary of how heatwaves change the outcomes of species interactions between the invader (*D. pandora*), the resident *Drosophila* (*D. pallidifrons*) and their shared parasitoids. (b) Summary of how the invading species changes the parasitoid-mediated coexistence between the dominant and inferior *Drosophila* species. Symbols of species are defined in Figure 2. Sizes of circles qualitatively represent the abundance of the adult populations. The thickness of the arrow represents the strength of the negative impact.

Similar demographic changes induced by extremes have been observed in studies of unmanipulated natural communities, facilitating the population growth of colonising species (Battisti, et al., 2006; Jiménez et al., 2011; Sorte, Fuller, & Bracken, 2010; Thibault & Brown, 2008; Wernberg et al., 2013). In comparison, species ranges and community compositions are sometimes stable despite substantial warming in mean temperature (Stuart-Smith, et al., 2010). Therefore, understanding the mechanisms (e.g., winter temperature, interspecific competition) preventing the establishment of species beyond their current range boundaries is key to predicting which climate factors will drive range shifts. High-resolution observational studies (Harley & Paine, 2009; Thibault & Brown, 2008) and experimental studies (Jentsch et al., 2007) like ours can be complementary to reveal the qualitative distinction between “press” (warming) and “pulse” (heatwave) climatic stresses in structuring biological communities (Easterling et al., 2000; Harris et al., 2018).

### Effects of climate-facilitated range-shifting species

Different speeds of range expansion and contraction lead to community reorganisation and novel species interactions (Parmesan et al., 1999; Sunday, Bates, & Dulvy, 2012). Novel interactions involving or driven by range-shifting species could become a crucial driver of community change (Descombes et al., 2020; Wallingford et al., 2020). Compared with typical biological invaders from other bioregions, climate-facilitated invaders from neighbouring communities are more likely to be sensitive to deviations from local temperatures, as they are already at the edge of their realized thermal niche.

In our study, the impact of the low-elevation specialist *D. pandora* on resident *Drosophila* community composition was only significant when the community also experienced heatwaves, consistent with our third hypothesis. Alexander et al. (2015) similarly found that novel competitive interactions were more influential when the abiotic conditions matched more closely those typical of the novel species’ source populations. Notably, the effects of invasion in our experiment were sometimes significant on individual species even when the introduction had not resulted in the successful establishment of *D. pandora*. For example, introducing this novel species significantly decreased the reproductive success of *Asobara* parasitoids and increased the abundance of *D. pallidifrons* under all thermal regimes. While the successful establishment of the novel species in the heatwave treatment indeed generated the largest shift in both the *Drosophila* and parasitoid communities, results on individual species suggest that the initial dispersal and subsequent interactions between neighbouring species and residents could also have a substantial influence on host-parasitoid dynamics.

### Important and unexpected indirect effects mediated by parasitism

We observed unexpected facilitation of *D. pallidifrons* by the invader, *D. pandora*, both in the short term (single-generation reproductive success) and long term (abundance in the census). *Asobara* parasitoids can develop successfully on monocultures of *D. pandora* (Nancie Bowley and Jinlin Chen, unpublished data). Therefore, the addition of *D. pandora* was expected to decrease the population size of *D. pallidifrons* in the long term through direct resource competition and apparent competition (Holt 1977). Instead, our regressions imply that introducing *D. pandora* lowers the efficiency of *Asobara* parasitizing *D. pallidifrons*. This effect was significant even when most offspring of the introduced *D. pandora* did not successfully develop into adults (in upland temperature or warming regimes). We suspect that the presence of *D. pandora* larvae spared *D. pallidifrons* larvae from attack by parasitoids (Figure 6a), a form of apparent facilitation or apparent mutualism (R. D. Holt & Lawton, 1994; Long, et al., 2012; Wilbur & Fauth, 1990). These parasitized *D. pandora* did not ultimately develop into adulthood due to competition, thus did not contribute to more parasitoid offspring. An additional possibility is that developing in *D. pandora* hosts leads to low-quality *Asobara* adults, perhaps because of the small body size or unsuitability of *D. pandora* its close relatives (i.e. *Drosophila bipectinata*, *D. pseudoananassae*) cannot be parasitized successfully by *Asobara* (Chia-Hua Lue, personal communications). If so, low-quality parasitoids would then have lower efficiency in parasitizing *D. pallidifrons*. Our example highlights that the consequences of novel host species integrating into host-parasitoid networks may not be correctly inferred on the basis of competition and parasitism studied in pairwise fashions. Experiments of how host competition alters parasitism success (in quantity and quality) will provide key insights.

Introducing *D. pandora* reduced the reproductive success of *D. pseudotakahashii*. Our analysis revealed that this was not a result of direct resource competition, but was also parasitoid-mediated. The reproductive successes of *D. pseudotakahashii* and *Asobara* parasitoids were associated positively. *Asobara* parasitoids cannot successfully develop on *D. pseudotakahashii* (Chia-Hua Lue, personal communication). However, parasitoids can alter patterns of competitive dominance among hosts (Cornell & Pimentel, 1978). A likely explanation for this observation is that parasitism pressure reduced competition between *D. pseudotakahashii* and other hosts that are susceptible to *Asobara*, such as *D. pallidifrons*, promoting coexistence between the dominant and inferior competitors (Figure 6b). Parasitoid performance is sensitive to both environmental temperatures and host species composition (Hance, et al., 2007; Thierry, Pardikes, Rosenbaum, et al., 2022; Thierry, Pardikes, Ximénez-Embún, et al., 2022). Top-town control could either stabilise (Ross et al. 2022; Sentis et al. 2013) or destabilise (Zarnetske et al. 2012) communities when the environment changes. In our experiment, parasitoids enhance the impact of the novel species, mediated by interaction modifications that were not expected *a priori*. The sensitivity of parasitoids implies that climate change and its related biotic changes pose a threat to the biodiversity maintained by their ecological functions.

### Limitations

Our manipulation of temperature and introduction treatments can reveal their overall causal effects. However, the underlying pathways and mechanisms are more speculative, based on analyses of associations and prior knowledge of species interactions. The novel species’ modification of the *Asobara*-*D. pallidifrons* interaction, and the modification by *Asobara* of *D. psudotakahashii*-*D. pallidifrons* competition need validation by direct experimental tests. Similar experiments that use communities with or without parasitoids will allow confirmation of how treatment effects are mediated by parasitoids.

While *Drosophila*-parasitoid communities provide tractable units for study at small spatial scales in the laboratory, our experimental setup inevitably differs in several respects from field conditions. One obvious difference is that parasitoids in our experiment are confined to a small space where it is easy for them to locate their hosts. This may explain why the larval parasitoid, *Asobara*, reached high abundance in our experimental communities. The ease of foraging might change competitive outcomes between our parasitoid species, and over-emphasize the mediating effects of *Asobara* parasitoids.

Nevertheless, our experiment reveals the potential for parasitoid-mediated effects. Besides, the situation of high parasitism rate has been seen in agricultural systems where parasitoids are constantly added as bio-control agents or even in some natural systems (Carton, et al., 1986).

### Practical implications for research, conservation and resource management

Our experiment shows that extreme events such as heatwaves can temporarily and strongly reduce the biotic resistance of local communities to invading species. This suggests that during and shortly after extreme climatic events may be the most crucial and effective time to facilitate the full recovery of the resident community (e.g., if endemic species within it are threatened by invasion), for example by manually removing invading species that are considered harmful (Wallingford et al., 2020). Our study also illustrates that parasitoids are sensitive to both heatwaves and novel hosts. Parasitoids are widely introduced to regions outside their natural ranges as biocontrol agents (Hardy et al., 1994). When selecting such biocontrol agents, applied entomologists may need to consider the interactions between changing climatic conditions and possible invading species, since unexpected interactions with novel species could inhibit the integration and effectiveness of biocontrol agents.

## Conclusion

The interdependence of abiotic and biotic drivers poses a major challenge to efforts to predict biological responses to climate change. Our work validates the role of heatwaves in facilitating range shifts, a pattern that is likely to occur more frequently in future alongside extreme climatic events. Constant warming and heatwave treatments had distinctive direct effects and associated biotic risks, emphasizing the need for more informed predictive research and targeting management practices against different components of climate change. Finally, parasitoids played a significant role in mediating the impacts of invasion; in particular, we found evidence for long-term facilitation between *Drosophila* hosts mediated by a larval parasitoid, a dynamic that has rarely been documented. Our work highlights the uncertainty of the ecological consequences of interactions between parasitoids and novel hosts under changing climates. It also underlines the importance of manipulative research in revealing the key species or interaction motifs that influence communities’ responses to climate change.

## Acknowledgements

We thank Jan Hrček (Czech Academy of Sciences) and Megan Higgie (James Cook University, Townsville) for their assistance in establishing laboratory cultures of *Drosophila* and parasitoids. Chia-Hua Lue provided information on the host-specificity of parasitoids, and Nancie Bowley contributed to a laboratory experiment testing parasitism of *D. pandora* by *Asobara* parasitoids. The Oxford Fly group (led by Ellie Bath and Irem Sepil) kindly shared facilities. Members of the Community Ecology Research Group, the Oxford Fly Group, and Jan Hrček’s Group provided advice on research plans and data analysis. We thank Chris Terry for his constructive advice on the manuscript.

