## Supplementary materials for "Experimental heatwaves and warming cause distinctive community responses through their interactions with a novel species"

**Supplementary Table 1. The effects of treatments and their interactions on the abundance of individual species.** Numbers of individuals species were log transformed for analysis. $P_{adjusted}$ are the adjusted p values following Bonferroni correction. Significant effects after correction for multiple comparisons are printed in bold.

| Focal species | Predictor | Coefficient | P value | $P_{adjusted}$ |
|---|---|---|---|---|
| *Asobara* sp. | **Intercept** | **6.23** | **<0.001** | **<0.001** |
| | **Warming** | **-0.33** | **<0.001** | **<0.001** |
| | **Heatwave** | **-0.40** | **<0.001** | **<0.001** |
| | **Introduction** | **-0.68** | **<0.001** | **<0.001** |
| | **Warming x Introduction** | **0.68** | **<0.001** | **<0.001** |
| | **Heatwave x Introduction** | **-0.59** | **<0.001** | **<0.001** |
| | **Block** | **-0.21** | **<0.001** | **<0.001** |
| *Trichopria* sp. | **Intercept** | **2.62** | **<0.001** | **<0.001** |
| | **Warming** | **-1.26** | **<0.001** | **<0.001** |
| | **Heatwave** | **-1.11** | **<0.001** | **<0.001** |
| | Introduction | 0.14 | 0.34 | 1 |
| | **Warming x Introduction** | **1.14** | **<0.001** | **0.001** |
| | Heatwave x Introduction | 0.16 | 0.57 | 1 |
| | **Block** | **0.47** | **<0.001** | **<0.001** |
| *D. pseudotakahashii* | **Intercept** | **4.21** | **<0.001** | **<0.001** |
| | **Warming** | **-0.72** | **<0.001** | **<0.001** |
| | **Heatwave** | **-2.27** | **<0.001** | **<0.001** |
| | **Introduction** | **-0.36** | **<0.001** | **0.004** |
| | Warming x Introduction | 0.13 | 0.39 | 1 |
| | Heatwave x Introduction | -0.001 | 0.99 | 1 |
| | **Block** | **-0.42** | **<0.001** | **<0.001** |
| *D. pallidifrons* | **Intercept** | **4.23** | **<0.001** | **<0.001** |
| | **Warming** | **-0.44** | **<0.001** | **<0.001** |
| | **Heatwave** | **0.42** | **<0.001** | **<0.001** |
| | **Introduction** | **0.66** | **<0.001** | **<0.001** |
| | Warming x Introduction | -0.29 | 0.003 | 0.14 |
| | Heatwave x Introduction | -0.05 | 0.54 | 1 |
| | **Block** | **0.46** | **<0.001** | **<0.001** |
| *D. birchii* | **Intercept** | **2.01** | **<0.001** | **<0.001** |
| | Warming | -0.25 | 0.32 | 1 |
| | Heatwave | -0.03 | 0.90 | 1 |
| | Introduction | -0.18 | 0.46 | 1 |
| | Warming x Introduction | 0.03 | 0.95 | 1 |
| | Heatwave x Introduction | -0.15 | 0.67 | 1 |
| | Block | -0.10 | 0.52 | 1 |
| *D. rubida* | **Intercept** | **3.03** | **<0.001** | **<0.001** |

| | | | | |
|---|---|---|---|---|
| | Warming | 0.32 | 0.01 | 0.37 |
| | Heatwave | 0.13 | 0.31 | 1 |
| | Introduction | 0.20 | 0.11 | 1 |
| | Warming x Introduction | -0.32 | 0.06 | 1 |
| | Heatwave x Introduction | -0.28 | 0.12 | 1 |
| | Block | 0.30 | <0.001 | 0.001 |

**Supplementary Figure 1. Temperature regimes.** (a) Recorded air temperatures in February 2017 at the high-elevation field site in Paluma, Queensland, Australia. The two curves coloured by red are days experiencing a heatwave. (b) Temperature treatments of the laboratory experiment. Colours indicate the weekly temperature of the corresponding temperature treatment (blue: 24°C, light (12h) and 18°C, dark (12h); orange: 25°C, light (12h) and 19°C, dark (12h); red: 30°C, light (12h) and 24°C, dark (12h) for five consecutive days of the specified week.)

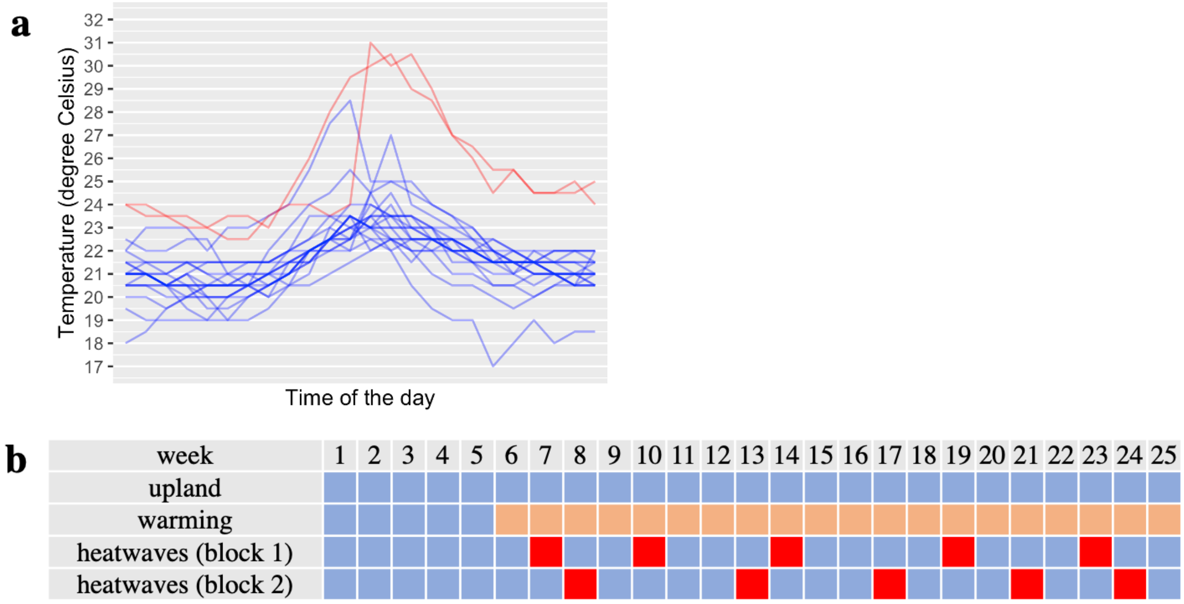

**Supplementary Figure 2. Censused abundance of individual species under temperature and introduction treatments.** (a) *Asobara* parasitoid; (b) *Trichopria* parasitoid; (c) *D. pseudotakahashii*; (d) *D. pallidifrons*; (e) *D. birchii*; (f) *D. rubida*. Each open symbol represents the adult abundance of the indicated species from one community replicate (n=5). Results of the post-hoc analyses are indicated by letters above the data. Groups labelled with different letters have statistically significant pairwise differences. The treatments are distinguished by colours as defined in the top panel. Shapes represent experimental block 1 (circle) and block 2 (triangle). The solid circles represent the means. Error bars show ±1 SD.

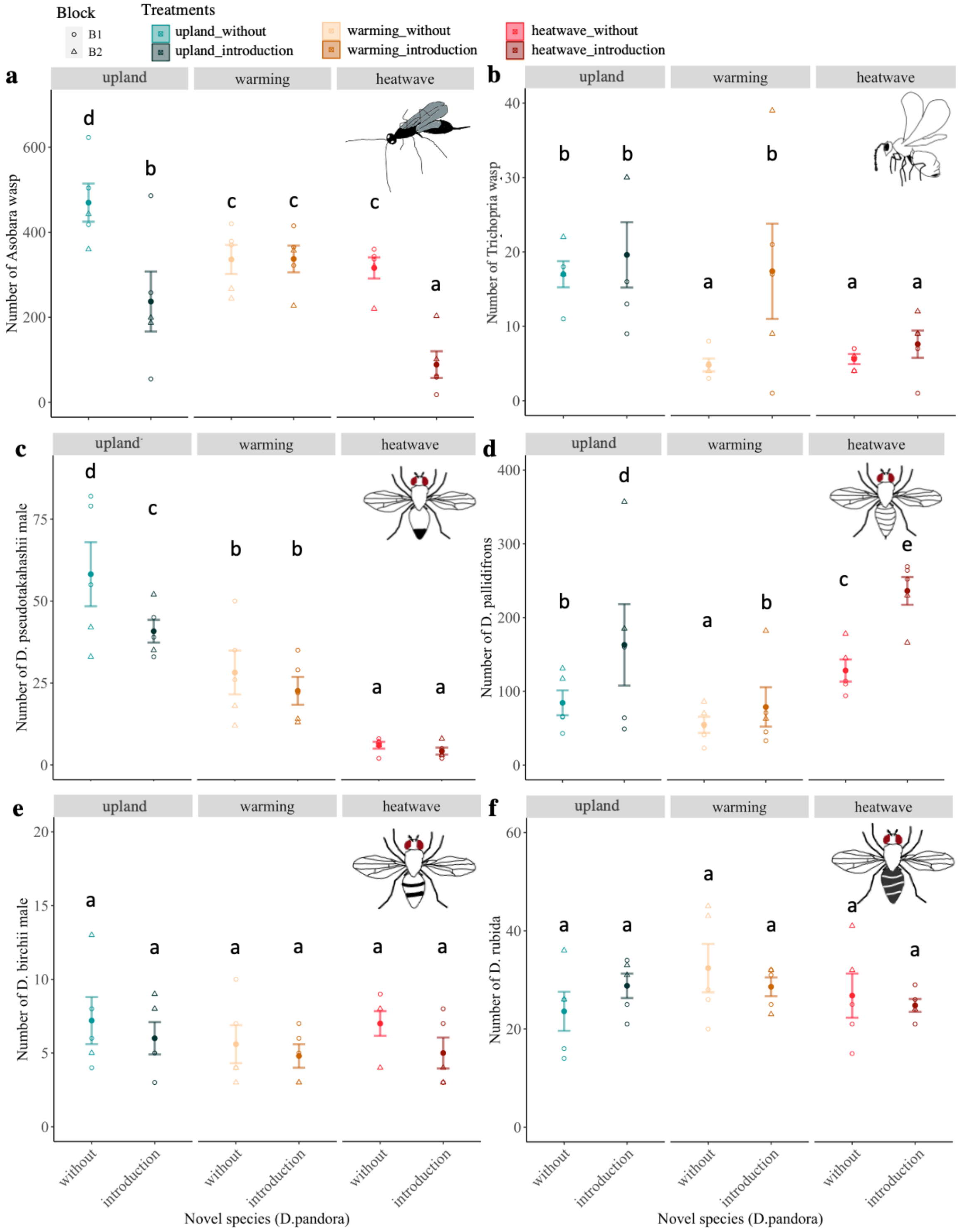

**Supplementary Figure 3. Reproductive success of individual species from tri-weekly sampling.** (a) *Asobara* parasitoid; (b) *Trichopria* parasitoid; (c) *D. pseudotakahashii*; (d) *D. pallidifrons*; (e) *D. rubida*; (f) *D. birchii*. Samples are viewed chronologically on the x axis. Temperature treatments are plotted in separate panel, and the *D. pandora* introduction treatment is indicated by the line type (dashed line: introduction; solid line: no introduction). Additionally, colours indicate the temperature and introduction treatment combination as defined in the top panel. Values for each cage are connected by a thin line. Thick lines represent mean values for the five replicate cages within each treatment combination.

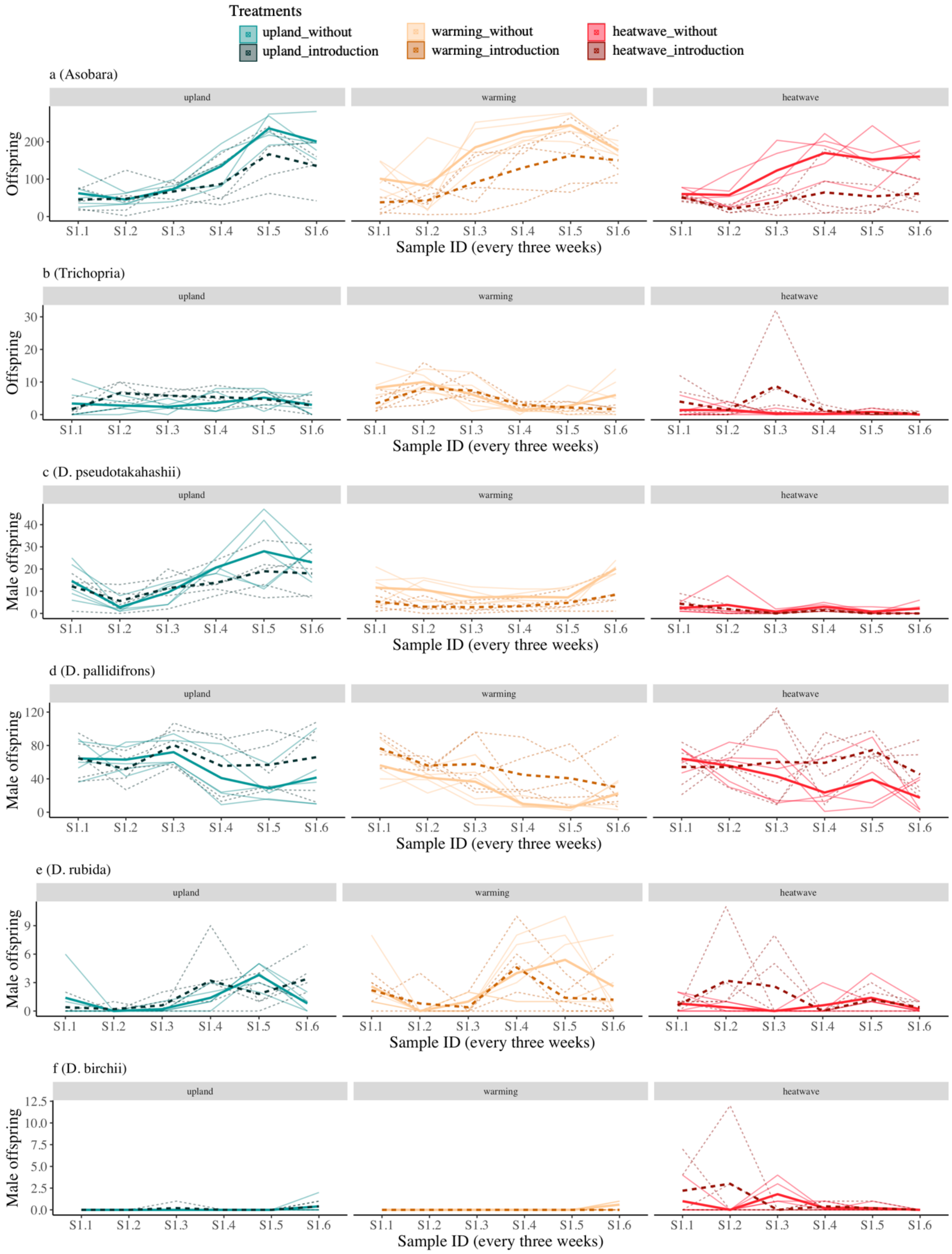
